# Finger Sweat Analysis Enables Short Interval Metabolic Biomonitoring in Humans

**DOI:** 10.1101/2020.11.06.369355

**Authors:** Julia Brunmair, Laura Niederstaetter, Benjamin Neuditschko, Andrea Bileck, Astrid Slany, Lukas Janker, Max Lennart Feuerstein, Clemens Langbauer, Mathias Gotsmy, Jürgen Zanghellini, Samuel M. Meier-Menches, Christopher Gerner

## Abstract

Metabolic biomonitoring in humans is typically based on the sampling of blood, plasma or urine. Although established in the clinical routine, these sampling procedures are often associated with a variety of compliance issues and are impractical for performing time-course studies. The analysis of the minute amounts of sweat sampled from the fingertip enables a solution to this challenge. Sweat sampling from the fingertip is non-invasive and robust and can be accomplished repeatedly by untrained personnel. This matrix represents a rich source for metabolomic phenotyping, which is exemplified by the detection of roughly 50’000 features per sample. Moreover, the determined limits of detection demonstrate that the ingestion of 200 μg of a xenobiotic may be sufficient for its detection in sweat from the fingertip. The feasibility of short interval sampling of sweat from the fingertips was confirmed in three time-course studies after coffee consumption or ingestion of a caffeine capsule, successfully monitoring all known caffeine metabolites. Fluctuations in the rate of sweat production were accounted for by mathematical modelling to reveal individual rates of caffeine uptake, metabolism and clearance. Biomonitoring using sweat from the fingertip has far reaching implications for personalised medical diagnostics and biomarker discovery.

## Introduction

Metabolomic studies seek to identify biomarkers for diagnosis, prognosis or therapy and hold great promise to improve clinical practice.^1^ Despite considerable progress with respect to the sensitive and parallel analysis of metabolites (*e.g.* by mass spectrometry^2,3^), the successful implementation of metabolites as biomarkers in the clinical setting still represents a major challenge.^4–6^ This is illustrated by strongly varying pharmacokinetics among individuals, which impacts significantly on drug responses.^7,8^ To the best of our knowledge, there is no analysis method available that would account for such individual responses. This may be partly due to the fact that study designs would need to include a large number of data points on a single individual. Most clinical research is based on the analysis of blood, plasma or urine,^9–11^ and due to compliance issues, these matrices do not allow for frequent sampling. Clearly, a non-invasive method is required to enable frequent sampling of the same individual in order to achieve individualised biomonitoring.

While fingerprints – the pattern of the ridge details left on a surface – have been used for the identification of individuals since the late 19^th^ century,^12^ their relevance for detecting metabolites, as well as drugs and their metabolites has only recently been discovered.^13,14^ While the drug substances detected in the fingerprint may originate from dermal contact or ingestion, the detection of drug-specific metabolites supports excretion from the sweat glands. Thus, we hypothesized that sweat from the skin surface may represent a promising source for metabolomic biomonitoring. Sweat is a hypotonic, slightly acidic biofluid secreted by the eccrine, apocrine and apoeccrine glands located on the skin surface.^15,16^ The composition of sweat comprises mainly water (∼99%), electrolytes, urea, lactate, amino acids and metal ions^17,18^ but also a variety of other endogenous metabolites, including peptides, organic acids, carbohydrates, lipids, lipid-derived metabolites as well as exogenous compounds (*e.g.* xenobiotics and drugs).^15,16,19,20^ Sweat composition is highly dynamic, changes significantly with pathological states and may reveal habits of diet, metabolic conditions or use of drugs and supplements.^11,18^ In fact, the analysis of sweat has already been reported to assess metabolic characteristics.^21,22^ Clinical assays based on the analysis of sweat exist and include the screening of newborn children for elevated chloride and sodium levels to confirm cystic fibrosis *via* pilocarpine stimulated iontophoresis or forensic and criminal investigations to test for illicit drug use.^11,16,23–25^ Furthermore, it has already been successfully demonstrated that the analysis of proteins contained in sweat enables not only the diagnosis of active tuberculosis but can also be used to screen for potential biomarkers for lung cancer,^10,26,27^ highlighting its potential for personalised medicine.^28^ Real-time monitoring of biomarkers was demonstrated with wearable sweat sensors for uric acid and tyrosine,^29^ interleukin-6 and cortisol^30^ or electrolytes such as sodium and ammonium ions and lactate.^31^

However, these studies typically assessed a small number of metabolites and relied on elaborate methods to collect sweat, including sweat patches or artificially forcing sweat production.^11,16,22^ This was necessary because the detection methods required relatively large absolute amounts of the metabolites. It is known that eccrine glands on the fingertips produce in the range of 50–500 nL cm^−2^ min^−1^ sweat.^32^ Thus, the analysis of metabolites from sweat of the fingertip may be achieved with sufficiently sensitive instrumentation.^33^ Sampling sweat from the fingertips for metabolomic biomonitoring requires no patient treatment or trained personnel and is safe and fast. In this study, we provide and benchmark an optimized workflow for the analysis of sweat from the fingertips. Proof-of-principle studies based on the consumption of coffee or caffeine capsules involving a high sampling rate provided evidence of the feasibility of the approach. Fluctuations in the rate of sweat production are accounted for by mathematical modelling of the conversion between xenobiotics and their metabolites, which supports personalised metabolomic biomonitoring in humans.

## Results

### Sweat from the fingertip is a rich source for metabolomic phenotyping

A straight-forward workflow was established for sampling sweat from the fingertip and processing the samples. In short, the hands are washed without soap and dried with a disposable paper towel prior to each sampling. For sweat collection, a sampling unit standardised to 1 cm^2^ diameter was then held between thumb and index finger for 1 min and was transferred with clean tweezers into an empty tube for storage (Figure 1A). The metabolites were extracted from the sampling units using aqueous conditions and the resulting solution was directly introduced into the liquid chromatography-mass spectrometry (LC-MS) system for analysis. Sample collection and processing required approximately 13 min per sample. Thus, sampling sweat from the fingertips is simple and fast. Sampling can be performed by untrained personnel in a highly frequent manner and the non-invasive nature of the sampling facilitates patient compliance. The data acquisition requires a further 7.5 min, which gives a total of approximately 20 min for the entire workflow per sample.

**Figure 1.**
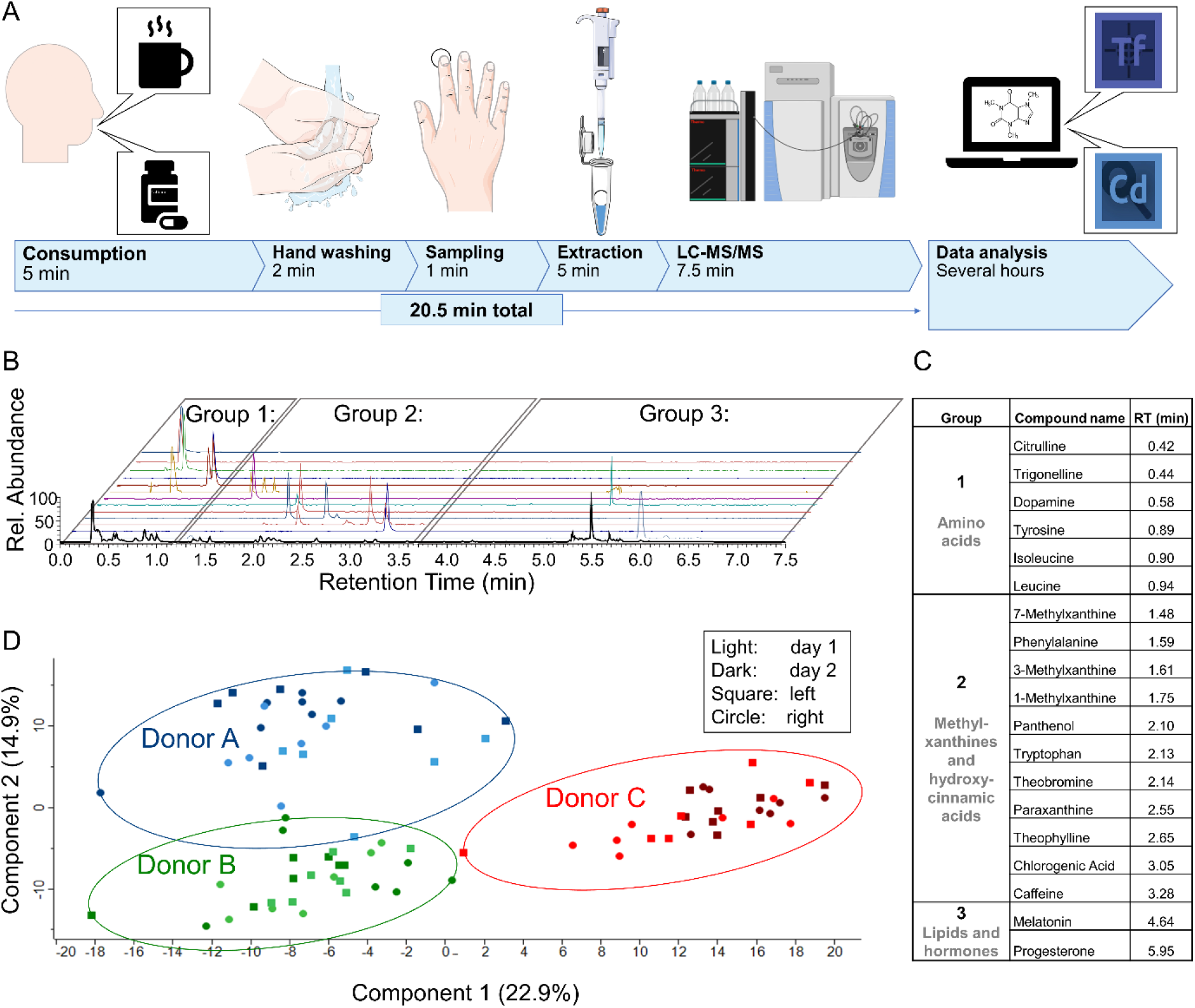
Sweat from the fingertips enables individualised biomonitoring based on metabolites. A straight-forward workflow for sweat sampling and processing was established and successfully applied to proof-of-principle studies to investigate caffeine metabolism in an individualised fashion. **(A)** Graphical summary of the workflow including coffee consumption, finger sweat sampling, the extraction of analytes and subsequent LC-MS/MS analysis as well as data analysis with respective working time in minutes. Tf, Tracefinder Software; Cd, Compound Discoverer Software (both Thermo Fisher Scientific) **(B)** Extracted ion chromatograms of key sweat components are shown. Based on their retention time, analytes were assigned to three groups. **(C)** Identities of sweat constituents depicted in Figure 1B. **(D)** A principle component analysis (PCA) of finger sweat samples derived from the left (square) and right (circle) hand of three donors is depicted before and after coffee consumption at two different days (light and dark colour). PCA successfully clustered the finger sweat samples according to the donors, suggesting individual finger sweat compositions.

**Table 1.**
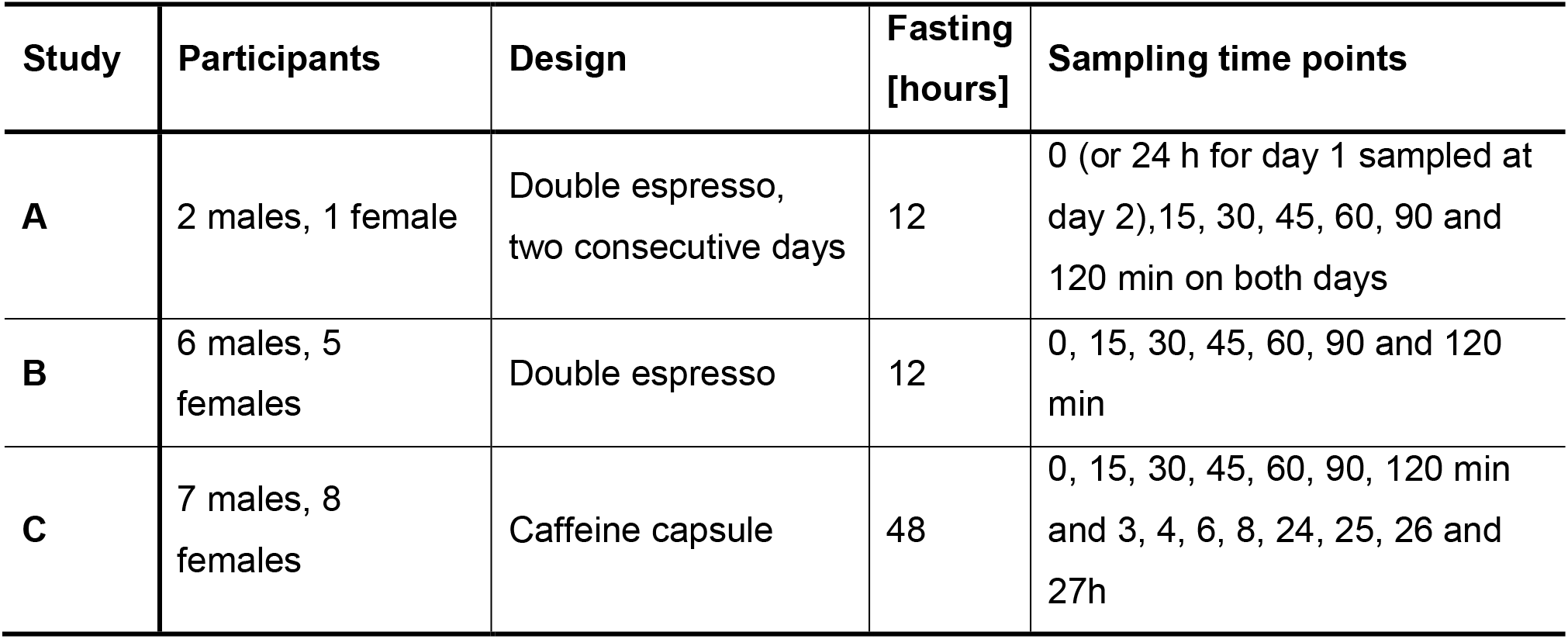
Overview of the three studies discussed in this publication.

Based on the known rates of sweat production in eccrine glands on the fingertips,^21,32^ the median sweat volume collected using this method can be estimated at around 100 nL per sample. Clearly, a sensitive method is required to detect and identify metabolites from such minute volumes. High-resolution MS using a Q Exactive HF orbitrap hyphenated with an ultra-high-performance liquid chromatography (UHPLC) system was used to perform untargeted metabolomics and typically delivered more than 50’000 reproducible sweat-specific features per analysis. Actually, many known as well as previously unknown metabolites were identified as sweat constituents with high confidence, including dopamine, progesterone and melatonin amongst others (Figure 1B–C). Subsequently, three volunteers were enrolled to participate in a coffee intervention study in order to evaluate the individual metabolomic composition and respective changes of sweat from the fingertip. Caffeine was chosen because of its widespread use as a central nervous system stimulant and its excellent oral bioavailability.^34,35^ The ingestion of an equivalent of a double espresso was already shown to affect sleep behavior^36,37^ and consequently, is expected to cause metabolic alterations. The volunteers collected sweat from the fingertips 7-times per day at different intervals on two consecutive days and using both hands (s. Methods, Study A). Principle component analysis of 250 compounds identified in sweat revealed that the samples clustered according to individuals (Figure 1C). This indicated that the sweat composition associated with a given individual dominated the variances derived from multiple sampling at different intervals and days. Moreover, we did not find notable differences of the sweat compositions between the left and right hand of an individual.

### Sampling sweat from the fingertip is reliable and robust

Biomolecules are characterised by LC-MS according to the retention time (RT), the accurate mass of the molecular ion derived from the full mass spectrum (MS1) and the fragmentation pattern determined by tandem mass spectrometry (MS2). The masses determined for 15 selected metabolites showed errors <2 ppm, which are typical for Q Exactive HF instruments (Table S1). Upon analysing 636 injections (s. methods, study A and C), the coefficient of variation (CV) of the RT determined for the internal standard caffeine-trimethyl-D9 was found to be 1% (Figure 2A). Caffeine-trimethyl-D9 was injected with every sample at 10 pg on column. The CV for the area under curve across all samples (n = 636) was 11%. The CV improved slightly when considering only the coffee study (study A, CV = 7%, n = 186) and remained similar when considering only the caffeine capsule (study C, CV = 10%, n = 450). This indicated that the performance of the LC-MS system was robust over the entire sample set. MS2 analysis provided clean spectra with high matching factors which supported the identification of metabolites previously known and unknown to be found in sweat, *e.g.* tryptophan^38^ and dopamine, respectively (Figure 2B). Such putative identifications were typically verified by external standards run under identical LC-MS conditions (Table S1). Caffeine and its three main metabolites paraxanthine, theobromine and theophylline were spiked onto sampling units in the range of 1– 100 pg μL^−1^. These samples were processed according to the above-mentioned procedures and linear calibration curves were obtained with associated R^2^> 0.997 (Figure 2C). At concentrations of 100 fg μL^−1^, these molecules were still detected with signal-to-noise ratios >100 on the Q Exactive HF. This results in an estimated limit of detection (LOD) for this method of around 10 μg L^−1^ sweat, depending on the ionisation efficiency of the given molecule. Based on this LOD and assuming a 20 L aqueous pool per body typically accessible for a xenobiotic *via* blood and lymph, the uptake of roughly 200 μg *in toto* of a stable substance may be sufficient for subsequent detection in the sweat from the fingertips. An extraction efficiency was calculated by comparing the 10 pg μL^−1^ spiked and processed standard to a directly injected 10 pg μL^−1^, yielding an extraction efficiency of 93%.

**Figure 2.**
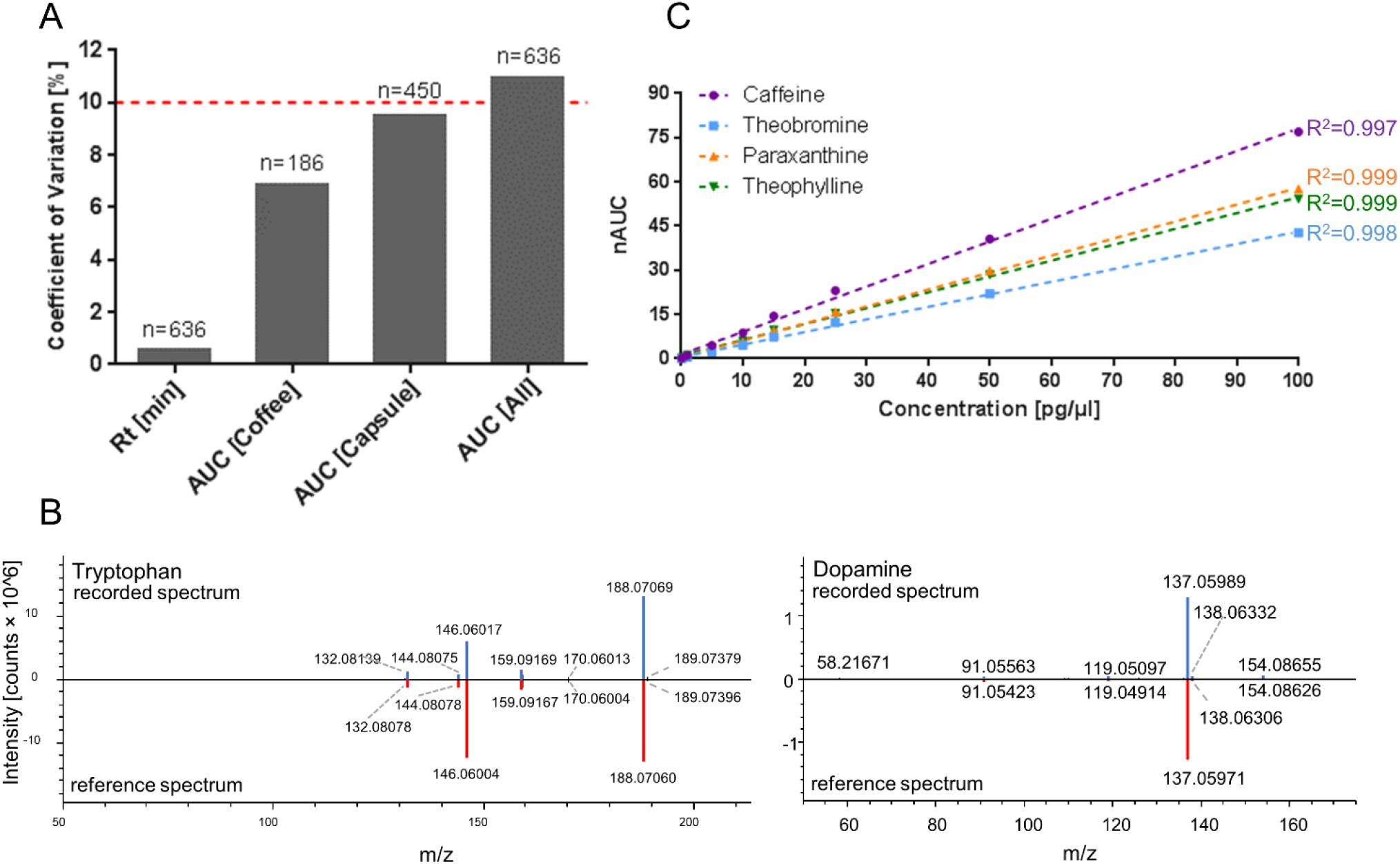
LC-MS/MS analysis of metabolites from sweat of the fingertips is precise and robust. **(A)** Coefficients of variation of the retention times and areas under the curve (AUC) for all LC-MS/MS runs, as well as AUCs for the coffee and caffeine capsule intervention studies were determined for the internal standard caffeine-trimethyl-D9. **(B)** Head-to-tail comparison of the recorded MS2 spectrum (blue) to the reference spectrum from *mzcloud* (red) of tryptophan (*left*) and dopamine (*right*) demonstrates high spectral quality supporting reliable compound identification. **(C)** Calibration curves for caffeine, theobromine, paraxanthine and theophylline with respective correlation factors (R^2^) are shown. All data is mean ± standard deviation. nAUC, normalised area under the curve.

### Coffee consumption revealed coffee-specific xenobiotics in sweat from the fingertips

As part of an intervention study, 11 participants consumed a standardised amount of coffee after a 12 h fasting period with regard to caffeine-containing food (s. Methods, study B). A sweat sample was collected before coffee consumption and subsequently after 15, 30, 45, 60, 90 and 120 min. Strikingly, the sweat from the fingertips 15 min post consumption revealed 35 coffee-specific xenobiotics (29%) of 121 metabolites presently identified by us from aqueous extracts of the roasted coffee beans used for this study, including among others caffeine, theobromine, theophylline, paraxanthine, methylxanthines, chlorogenic acid, trigonelline, methylsuccinic acid, quinic acid and iditol (Supplementary Table). The areas under the curve (AUCs) increased significantly in all donors as early as 15 min after coffee consumption, as demonstrated for caffeine, chlorogenic acid and trigonelline (Figure 3A). The time-dependent sampling revealed differences in pharmacokinetic properties of the coffee-specific xenobiotics detected in sweat, especially regarding uptake and clearance rates. For example, the AUCs of caffeine and chlorogenic acid peaked after 15 min, followed by rapid clearance, while the AUCs of the dimethylxanthines increased steadily over time (Figure 3B). Several coffee-specific metabolites displayed a number of isomers in their extracted ion chromatograms*, e.g.* chlorogenic acid (*m/z* 355.1024, RT = 3.05 min) showed at least five isomers as verified on MS2 level (Figure 3C). The ratio of the relative peak intensities of chlorogenic acid and its isomers was conserved when comparing coffee extracts and sweat from the fingertip. This indicated a rather efficient uptake of these metabolites into the water-soluble body compartment and delivery into sweat glands without specific discrimination. Chlorogenic acids and its isomers were not observed prior to coffee consumption. Such a comparative analysis strategy may be used to screen for other coffee-specific xenobiotics distributed to sweat glands in a systemic fashion and may even be extended to the feature level, *e.g.* the yet unidentified feature with *m/z* 337.0920 (Figure S1). These findings provide evidence that ingested xenobiotics may be robustly detected in the sweat from the fingertip, which mirrors their pharmacokinetic properties. While chlorogenic acid was not detected before coffee consumption and increased significantly shortly after, the abundance of the dimethylxanthines increased more slowly on top of an existing pool.

**Figure 3.**
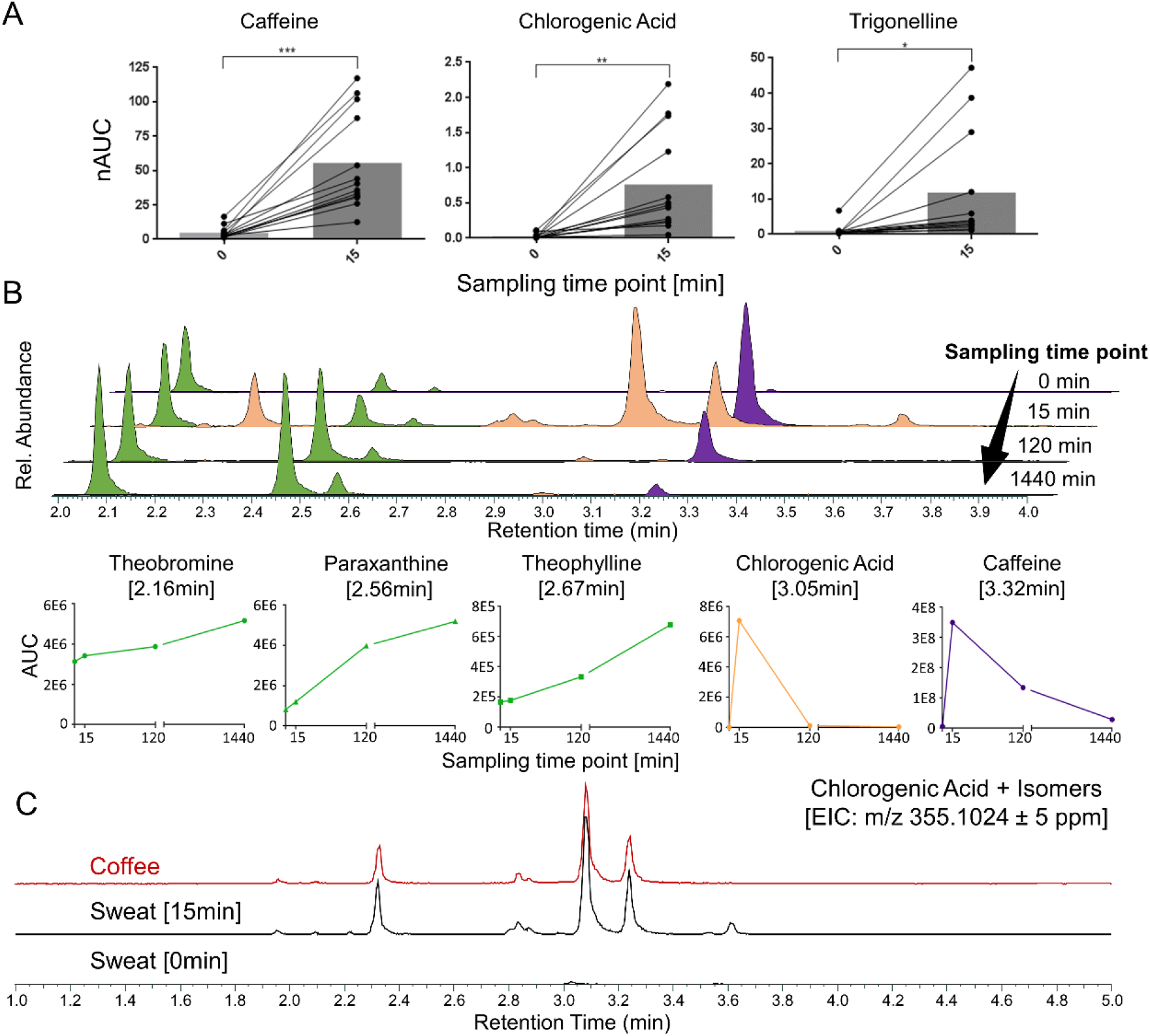
Xenobiotics are detected in a time-dependent manner in sweat from the fingertips after coffee consumption. **(A)** Levels of normalized areas under the curve (nAUCs) for caffeine, chlorogenic acid and trigonelline, before (0) and 15 min (15) after coffee consumption are shown, demonstrating a significant increase in all individuals after 15 min. Paired *t*-test performed for 13 (*df* = 12) donors delivered the following results: for caffeine *t* = 5.531, *p*-value = 0.0001; for chlorogenic acid *t* = 3.072; *p*-value = 0.0030; for trigonelline *t* = 2.657, *p*-value = 0.0209. **(B)** Characteristic kinetics of different compounds are depicted for one donor (Donor 3, Study A). 1440 min is the sampling time point before consumption on the second sampling day. Whereas caffeine and chlorogenic acid were found to rise quickly after coffee consumption followed by rapid clearance, the levels of theobromine, paraxanthine and theophylline increased more slowly within the observation period. (**C**) Similarity of extracted ion chromatograms (EIC) of chlorogenic acid and its isobars from coffee extracts and from sweat of the fingertips 15 min after coffee consumption. The corresponding sample collected just before coffee consumption (0 min) served as negative control.

### Analysis of sweat from the fingertips supports the determination of individual metabolic traits

The metabolism of caffeine by different enzymes is well known,^39^ resulting in the formation of several metabolic products, all of which were successfully identified by us in sweat from the finger tips (Figure 4A, Table S1). Since the metabolites of caffeine featured time-dependent profiles upon coffee consumption, we wondered whether the analysis of sweat from the fingertips supported the characterisation of individual traits of caffeine metabolism. To that aim, 15 volunteers refrained from consuming caffeine-containing products for 48 h before consuming a single 200 mg caffeine capsule. Again, sweat from the fingertips was sampled repeatedly over 27 h (s. Methods, study C). Six volunteers participated in the coffee consumption study (study B) and the caffeine capsule study (study C). For these individuals, the longer fasting time improved the baseline and revealed a significant decrease of dimethylxanthine levels (Figure 4B). Ingestion of the caffeine capsule significantly increased the abundance of caffeine in sweat from the fingertips in all donors already after 15 mins. The caffeine abundance remained elevated for at least 480 min in all donors, but returned close to baseline after 24 h. The abundance of paraxanthine increased more slowly and peaked between 360–480 min post ingestion (Figure S2). Similarities in the response of all volunteers to the caffeine capsule were revealed by statistical analysis of the untargeted sweat samples with volcano plots, which identified 54 significantly up-regulated features within 360 min post-ingestion, including caffeine, theophylline and paraxanthine (Figure 4C). Theophylline and paraxanthine reflected metabolic activity within each volunteer. After 24 h post-ingestion, 47 features were significantly upregulated, including caffeine and dopamine. Importantly, dopamine is an endogenous substance, which is not directly related to caffeine metabolism. Despite these similarities across all volunteers, rather striking differences regarding caffeine metabolism were discerned when considering individual metabolic time-courses (Figure 4D). For example, donor 1 displayed a sharp increase in caffeine abundance, which remained rather constant over 480 min, while paraxanthine abundance increased steadily during this time period. In contrast, donor 2 featured also an increase in caffeine abundance, but started with a higher theobromine baseline, which also represented the main metabolite. These findings suggest that sampling sweat from the fingertip is also of interest for characterising personalised metabolic traits and individual responses to xenobiotic interventions.

**Figure 4.**
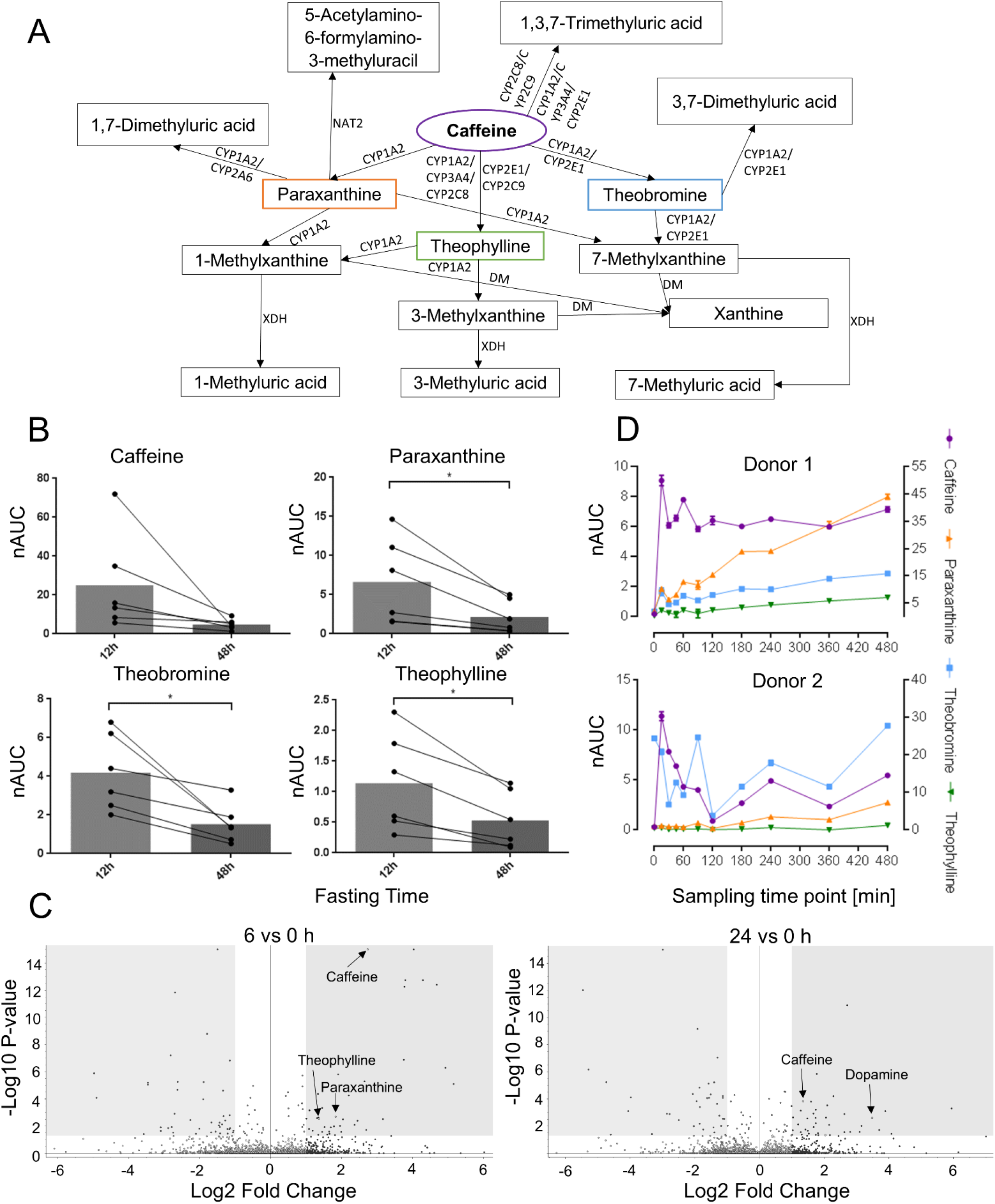
Consumption of a caffeine capsule enables individualised metabolic profiling from sweat of the fingertips. **(A)** Caffeine metabolism including known metabolic routes, metabolites and related enzymes: CYP, Cytochromes P450; NAT2, N-acetyltransferase 2; XDH, Xanthine dehydrogenase; DM, demethylase. These metabolites were all detected in sweat from the fingertip. **(B)** Differences between 12 h *vs* 48 h of fasting caffeinated foods and drinks for 6 individuals participating in the coffee as well as in the caffeine capsule studies. Longer fasting significantly reduced the amounts of xenobiotics in sweat from the fingertips. Paired *t*-test (6 donors x 2 time points) for caffeine (*p*-value = 0.1033, *t* = 51.990, *df* = 5), paraxanthine (*p*-value = 0.0297, *t* = 3.012, *df* = 5), theobromine (*p*-value = 0.0203, *t* =3.353, *df* = 5) and theophylline (*p*-value = 0.0118, *t* = 3.866, *df* = 5). nAUC = normalised area under curve. **(C)** Volcano plots illustrate the similarities of metabolite regulations in 15 individuals 6 h and 24 h after the intake of a caffeine capsule. For caffeine the *p*-value is 1E-15, for theophylline 0.0025 and for paraxanthine 0.0021 in case of 6 vs 0 h and for caffeine the p-value is 0.0002 and for dopamine 0.0025 corresponding to the 24 h vs 0 h. **(D)** Exemplary metabolic profiles of two donors, demonstrating individual differences in metabolic properties regarding caffeine metabolism as exemplified by preferential formation of paraxanthine in donor 1 in contrast to theobromine in case of donor 2. Caffeine is displayed on the right y-axis, while theobromine, paraxanthine and theophylline are displayed on the left y-axis. nAUC, normalised area under the curve. All data is mean ± standard deviation.

### Mathematical modelling of metabolic time-course measurements is successfully implemented to quantify individual metabolic traits

Fluctuations in the rate of sweat excretion cause significant variance in the collected sweat volumes. This represents a fundamental challenge for the time-course analysis of sweat from the fingertips. For example, the apparent down regulation of all analytes at 120 min in donor 2 (Figure 4C) strongly suggests that at that time point less sweat was collected in comparison to the adjacent measurements (s. Figure 5B, arrow). Moreover, the magnitude of this effect on the apparent concentration is unknown. We used dynamic metabolic network modelling to discern the effects of the sweat volume on the measured time series of caffeine degradation in the body (s. Methods). In brief, caffeine uptake and clearance *via* its major metabolic products paraxanthine, theobromine and theophylline can be described by first order kinetics (Figure 5A).^40,41^Additionally, we assumed that the sweat volume is constant for all metabolites at the same time point and is described as a function of time. The assumption holds if the modelled metabolites are not reabsorbed during sweating.

**Figure 5:**
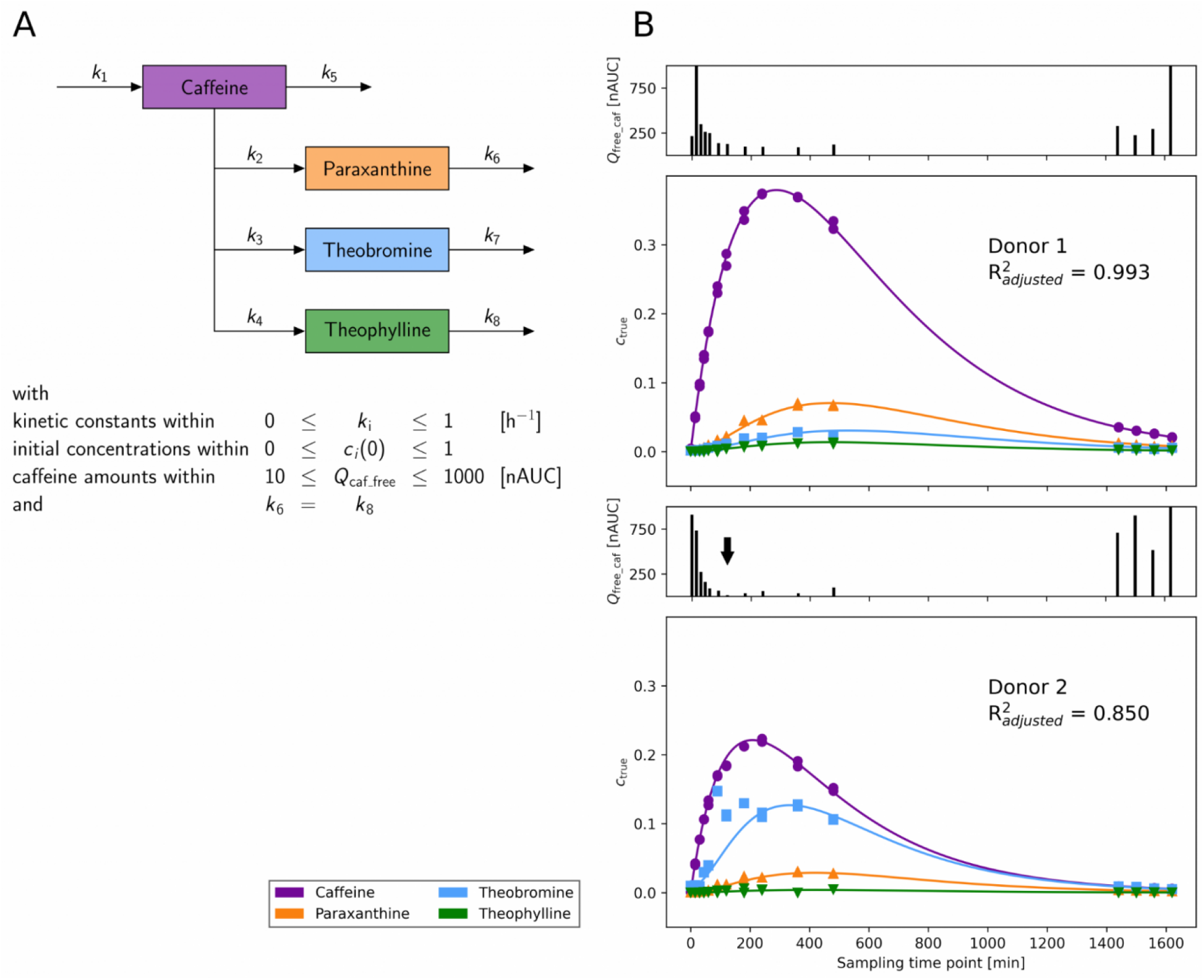
Metabolic networks facilitate the discovery of personalised kinetic parameters. **(A)** Network of caffeine and its major metabolites that was used for fitting of the concentration time series. **(B)** Big panels show the fitted concentration time series of caffeine, paraxanthine, theobromine and theophylline for donors 1 and 2 (compare Figure 4C). The lines refer to the fitted concentration and the symbols refer to the measured values divided by the fitted caffeine amount (*Q*_free_caf_) at each time point. Small panels show the size of the estimated *Q*_free_caf_, which consists of time-independent constants and time-dependent the sweat volumes. The arrow marks the (small) sweat volume of donor 2 at 120 min (see text). A visual representation of the caffeine amount on the fit is shown in Figure S3.

The resulting mathematical model was fitted to each donor and recovered the kinetic constants, the initial concentrations of paraxanthine, theobromine and theophylline and the sweat volumes at each time point, as exemplified for donors 1 and 2 (Figure 5B and Supplementary Table S2). In both cases our model accurately described individual caffeine metabolism. Thus, individual metabolic traits can be obtained from the shape of the curves.

Interestingly, the kinetic constants for conversion of caffeine into its metabolic products (*k*_2_, *k*_3_, *k*_4_) are close to literature values for blood plasma, however, the absorption constant *k*1 is considerably reduced in sweat (Supplementary Table S2).^42^ Moreover, our method supports the identification of individual traits in metabolism. Whereas the fractional conversion of caffeine to the main metabolic product paraxanthine in donor 1 is similar to what is described as population average^42,43^ we saw substantial differences for donor 2, who displayed theobromine as the main metabolic product of caffeine (Table S3).

### Targeted assays can be established from untargeted metabolomics for clinical implementation

The described untargeted metabolomics approach represents a powerful discovery tool for metabolic biomonitoring. In order to evaluate the feasibility of clinical implementation, we established a targeted assay for caffeine, and the primary metabolites theobromine, theophylline and paraxanthine on a triple quadrupole MS using multiple reaction monitoring (MRM, s. Supplementary Information and Table S4). For this purpose, five volunteers consumed a standardised coffee on three independent days after a 12 h caffeine-free fasting period and samples were collected at different time intervals in analogy to study B. The assay was validated and revealed linear ranges between 0.25–150 pg of the respective metabolites on column (Figure S5). LOD values were determined around 0.2 pg per collected sweat sample. The overall process efficiencies were generally >88% and the precision of 50 pg spiked metabolite was <2% (Table S5), while the overall CV of the AUC of caffeine 5 h after coffee consumption of all volunteers over three independent days was 22% (Figure S6). This suggest that targeted assays based on the analysis of sweat from the fingertips can be successfully established directly from metabolomic discovery studies.

## Discussion

The present study provides evidence that untargeted metabolomics of sweat from the fingertips is a powerful approach for human biomonitoring by supporting patient compliance. The sample collection is non-invasive and can be accomplished by untrained personnel. Time-course analyses with frequent sampling can be performed due to the facile collection procedure. Sample processing and analysis can be accomplished in 20 min per sample, which is desirable for large scale metabolomic studies.

Sweat from the fingertip represents a rich source for metabolic biomonitoring. Considering that a given metabolite may be represented in an LC-MS experiment by several features due to different adducts and charge states,^44^ it may be concluded that several thousand distinct metabolites can be detected and identified in sweat from the fingertip using this methodology. The analysis is robust and sensitive with limits of detection of metabolites typically found in the sub-picogram range per fingertip. Indeed, detection limits in sweat from the fingertip were found far below the values observed by us previously using other methodologies.^45^ As a result, numerous endogenous metabolites were identified with this method, which have not yet been described in sweat, such as secondary metabolites, including dopamine, progesterone and melatonin (Figure 1). This highlights the potential of this approach to successfully identify low-abundant metabolites, which are challenging to detect in other biofluids (*e.g.* melatonin in blood or plasma).^46,47^ Analysis of the area under the curve of the internal standard revealed an overall coefficient of variation of 11% across 636 samples and indicated acceptable precision (Figure 2). Furthermore, extrapolation of the limits of detection suggest that the uptake of xenobiotics in sub-milligram amounts by an individual may be evidenced in sweat from the fingertip. This finding has broad implications for medical diagnostics, since this strategy may be adapted for monitoring whether patients actually take prescribed pharmaceuticals. Moreover, this approach may be promising for nutritional supervision as highlighted by the detection of many caffeine metabolites after coffee consumption.

Proof-of-principle intervention studies were successfully carried out and support the applicability of the method. In three separate studies, volunteers were asked to consume a standardised coffee or ingest a caffeine capsule after a caffeine– and theobromine-free diet for 12 or 48 h. After consumption or ingestion, sweat samples were collected in short time intervals. Consuming a coffee led to a significant upregulation of caffeine, chlorogenic acid and trigonelline as quickly as 15 min after ingestion in all volunteers (Figure 3). This suggested a fast uptake and distribution of these xenobiotics, which was further underlined by largely overlapping isobar profiles of coffee extracts and sweat from the fingertip (Figure 3C). Altogether, 35 coffee-specific metabolites were detected in sweat from the fingertip.

In addition to the detection of xenobiotics in sweat from the fingertip, these compounds may be metabolised in the body, with metabolic activities subjected to individual variation. Thus, volunteers ingested a caffeine capsule and indeed, all known caffeine metabolites were successfully identified in sweat from the fingertip. Interestingly, statistical analysis of the untargeted metabolomics data from this study revealed a significant upregulation of caffeine and of the metabolic products theophylline and paraxanthine across all volunteers after 480 min (Figure 4). After 24 h, the volunteers featured significantly increased levels of dopamine with respect to 0 min. Being an endogenous metabolite, it is plausible to assume this upregulation corresponded to a physiological response of caffeine ingestion. Increased dopamine levels were already observed upon caffeine consumption.^48^ Consequently, sweat from the fingertips may not only reveal ingested xenobiotics, but also endogenous metabolic products of a physiological response to bioactive molecules. We have previously described individual opposing responses with regard to anti-inflammatory effects after coffee consumption.^49^ Such studies required the collection of blood from volunteers and this could be facilitated by analysing sweat from the fingertip.

Individual metabolic traits were then observed by analysing time-dependent metabolite profiles upon ingestion of the caffeine capsule (Figure 4D). We found that sweat from the fingertip may be successfully used for the personalised assessment of such metabolic activities. Importantly, this strategy may be extended to other xenobiotics or drugs and their causally related metabolic products in order to obtain insight into specific processes of human metabolism in an individualised manner. Variations in the sweat volume over the course of the study represent a challenge for normalisation and data interpretation. Mathematical modelling overcame this issue by considering numerous molecular constraints such as substrate-product relations. Successful modelling has two central prerequisites: firstly, the measurement of at least two metabolites with known dynamics and, secondly, no reabsorption of said metabolites during sweating. This allows us to compute a sweat volume that is proportional to all metabolites at each time point. This approach was capable of delivering estimates of individual rate constants for drug uptake, metabolism and clearance (Figure 5). Sampling sweat from the fingertip enables time-course studies, which are evaluated by means of conversion rates of metabolically related substance classes and thus, circumvent the requirement of absolute quantitative information of a single measurement. Further research is currently performed in order to consolidate the potential of sampling sweat from the fingertips, which may lead to the development of novel clinically-relevant assays.

In summary, we present a powerful method for metabolomic biomonitoring in humans using sweat from the fingertips, which offers numerous practical applications for medical diagnosis, including the detection of xenobiotics or metabolic traits. Although this approach centres on metabolomics discovery studies using dedicated high-resolution instrumentation, we demonstrated the successful transfer to a targeted assay, which may be quickly implemented in clinical laboratories.

## Methods

### Reagents and Chemicals

LC-MS grade methanol, water, acetonitrile and formic acid used during sample preparation and LC-MS/MS analysis were purchased from VWR chemicals (Vienna, AT). Xenobiotic and metabolite standards (caffeine, theobromine, theophylline, paraxanthine, 1-methylxanthine, 3-methylxanthine, 7-methylxanthine, 1-methyluric acid, 3-methyluric acid, 1,7-dimethyluric acid, 3,7-dimethyluric acid and 1,3,7-trimethyluric acid, chlorogenic acid, xanthine, 5-Acetylamino-6-formylamino-3-methyluracil, dopamine and proteinogenic amino acids) were either purchased from Sigma Aldrich (Vienna, AT) or Honeywell Fluka (GER). Caffeine capsules were bought from Mach dich wach! GmbH (Germany). Sampling units were made from filter papers (Kimtech Science) using a circular puncher of 1cm^2^.

### Standard Solutions and Calibration Samples

Stock solutions of 1 mg mL^−1^ of the analytical standards and the internal deuterated standards caffeine-trimethyl-D9 and N-acetyl-tryptophan in methanol were prepared and stored at 4 °C. For caffeine, paraxanthine, theobromine and theophylline calibration curves were generated by spiking onto sampling units with the following concentrations: 0.1, 1, 5, 10, 15, 25, 50, and 100 pg μL^−1^. The internal deuterated standards were prepared at a concentration of 1 pg μL^−1^ in an aqueous solution containing 0.2% formic acid, which served as the extraction solution for all samples.

### Cohort Design

Altogether, 7 males and 8 females with ages between 20–50 years were enrolled in the studies after providing written informed consent. These experiments were approved by the ethical committee of the University of Vienna (no. 00337). Volunteers had different dietary habits regarding the consumption of coffee; rare to regular consumption. Prior sampling, subjects were asked to fast caffeinated food (*e.g.* chocolate) and drinks (*e.g.* coffee, tea, and energy drinks) for a period of 12 h or 48 h. Afterwards, finger sweat samples were collected just before volunteers consumed a coffee (equivalent to a double espresso) or a 200 mg caffeine capsule (0 min). Finger sweat samples were generally collected after 15, 30, 45, 60, 90 and 120 min in case of the consumption of coffee. Additional samples were collected after 3, 4, 6, 8, 24, 25, 26 and 27 h after ingestion of the caffeine capsule. Volunteers may have participated in more than one study. It was ensured that the volunteers did not touch the prepared coffee with their fingers.

### Collection of Sweat from the Fingertip

Sampling units of 1 cm^2^ circular surface were pre-wetted with 3 μL water and provided in 0.5 mL Eppendorf tubes. For each sweat collection, volunteers cleaned their hands using warm tap water and dried them with disposable paper towels. Donors kept their hands open in the air at room temperature for 1 min. Then, the sampling unit was placed between thumb and index finger using a clean tweezer and held for 1 min. Sweat formation was not forced. Filters were transferred back to labelled 0.5 mL Eppendorf tubes using a clean tweezer and stored at 4 °C until sample preparation.

### Sample Preparation

Coffee extracts were prepared taking an aliquot of 1 mL of a 250 mL coffee cup used for study A and B, which was centrifuged for 10 min at 15000 rcf. The supernatant was diluted 1: 100, 1: 1000 and 1: 10000 with the extraction solution consisting of an aqueous solution of caffeine-trimethyl-D9 (1 pg μL^−1^) with 0.2% formic acid. The dilutions were again centrifuged before analysis by LC-MS/MS.

For the extraction of metabolites from the sampling units, 120 μL of the extraction solution consisting of an aqueous solution of caffeine-trimethyl-D9 (1 pg μL^−1^) with 0.2% formic acid was added into the 0.5 mL Eppendorf tube containing the sampling unit. The metabolites were extracted by pipetting up and down 15 times. The sampling unit was pelleted on the bottom of the tube and the supernatant was transferred into HPLC vials equipped with a 200 μL V-shape glass insert (both Macherey-Nagel GmbH & Co.KG) and analysed by LC-MS/MS. Additionally, unused sample units were extracted similarly to determine their metabolite background and potential contaminants.

### LC-MS/MS Analysis

A Q Exactive HF (Thermo Fisher Scientific) mass spectrometer coupled to a Vanquish UHPLC System (Thermo Fisher Scientific) was employed for this study. Chromatography was performed using a Kinetex XB-C18 column (100 Å, 2.6 μm, 100 × 2.1 mm, Phenomenex Inc.). Mobile phase A consisted of water with 0.2% formic acid, mobile phase B of methanol with 0.2% formic acid and the following gradient program was run: 1–5% B in 0.3 min and then 5–40% B from 0.3– 4.5 min, followed by a column washing phase of 1.4 min at 80% B and a re-equilibration phase of min at 1% B resulting in a total runtime of 7.5 min. Flow rate was set to 500 μL min^−1^, the column temperature to 40 °C, the injection volume was 10 μL and the injection peak was found at RT = 0.3 min. All samples were analysed in technical replicates. An untargeted mass spectrometric approach was applied for compound identification. Electrospray ionisation was performed in positive and negative ionisation mode. MS scan range was *m/z* 100–1000 and the resolution was set to 60000 (at *m/z* 200). The four most abundant ions of the full scan were selected for HCD fragmentation applying 30 eV collision energy. Fragments were analysed at a resolution of 15000 (at *m/z* 200). Dynamic exclusion was applied for 6 s. The instrument was controlled using Xcalibur software (Thermo Fisher Scientific).

### Data Analysis

Raw files generated by the Q Exactive HF instrument were analysed using the Compound Discoverer Software 3.1 (Thermo Fisher Scientific). Identified compounds were manually reviewed using Xcalibur 4.0 Qual browser (Thermo Fisher Scientific) and the obtained MS2 spectra were compared to reference spectra, which were retrieved from *mzcloud* (Copyright © 2013–2020 HighChem LLC, Slovakia). The match factor cut-off from *mzcould* was 80, while the mass tolerances were 5 and 10 ppm on MS1 and MS2 levels, respectively. Moreover, the identity of compounds suggested by Compound Discoverer was verified by analysing purchased standards using the same LC-MS method. The Tracefinder Software 4.1 (Thermo Fisher Scientific) was used for peak integration and calculation of peak areas. The generated batch table was exported and further processed with Microsoft Excel, GraphPad Prism and the Perseus software,^50^ the letter being used for the principle component analysis.

### Statistical Analysis

Two-tailed, paired t-tests were performed for mass spectrometry data using GraphPad Prism (Version 6.07) to evaluate the significance of the abundance increase/decrease of compounds and their metabolites. Volcano plots were obtained using Compound Discoverer Software, setting the p-value for statistical significance to 0.05 and the log_2_ fold change to 1.

### Mathematical Modelling

A mathematical formulation of the problem of fluctuating sweat volumes is given in Equation 1, where *M*_apparent_ is the measured amount of a metabolite and *C*_true_ is the true concentration. *SV* is a time-dependent factor that represents the sampled sweat volume. The following assumptions are made in setting up this model:

- caffeine metabolism can be described by mass-action kinetics in a one-compartment body model^40,41^
- distribution volumes are constant and identical for caffeine, paraxanthine, theobromine, and theophylline^42^
- apparent metabolite concentrations are proportional to the sweat volume (s. Figure S4)
- sweat volumes are time-dependent.

The resulting mathematical model is given in the Supplementary Information, Equations S1-S5. Its solution can be given analytically as listed in the Supplementary Information, Equations S6– S9. In all equations the upper-case constants *M*, *C*, *V* have their respective units, whereas their lower-case equivalents are normalised and thus unit-less. This leads to the fact that a unit-less concentration term is used for fitting and the mass-unit is introduced by the amount of free caffeine *Q*_free_caf_(*t*) (Equation 2), which not only represents the sweat volume *SV* but also the dose of ingested caffeine (200 mg) and two unknown factors (bioavailability and distribution volume). As opposed to traditional pharmacokinetic models,^51^ this model did not explicitly include said parameters since they are factors analogous to *SV*, which cannot all be determined unambiguously. This means that in this model only the caffeine amount denoted as *Q*_free_caf_(*t*) was fitted, however it includes all the above-mentioned terms and because of this has the mass-unit nAUC (Equation 2). Nevertheless, two points should be kept in mind, firstly, *Q*_free_caf_(*t*) scales the fitted curve on the y-axis, whereas its shape – and thus the kinetic parameters that describe individual metabolic differences – can be unambiguously determined and, secondly, all constants except for SV in Q_free_caf_(*t*) are time-independent and therefore the variance of Qfree_caf(*t*) over time solely depends on the variance of SV.

Note that since *Q*_free_caf_(*t*) is not constant over time and a fitting parameter at each instance, the amount of data needed for fitting all parameters is equal to the number of time points plus the number of parameters of the kinetic model. This requires the simultaneous fitting of the dynamics of multiple metabolites upon assuming that at each time point *Q*_free_caf_(*t*) is constant for similar metabolites. By doing so the number of data points that can be fitted is multiplied, whereas the number of parameters for *Q*_free_caf_(*t*) stays constant, and thus enables data fitting.

Caffeine and its major metabolic products paraxanthine, theobromine and theophylline were modelled subject to the following constraints: first order kinetics for all reactions (*k*_1_ to *k*_8_); initial unit-less concentration of 0 for caffeine and 0 ≤ *c_i_*(0) ≤ 1 for other metabolites; variability of caffeine amount in two orders of magnitude 10 nAUC ≤ *Q*_free_caf_(*t*) ≤ 1000 nAUC and equal elimination parameters of paraxanthine and theophylline (*k*_6_ = *k*_8_) since they have the biggest influence on the time of maximum concentration, which was reported to be similar for both metabolites.^52^ The system of equations denoted in Equation (2) combined with Supplementary Information, Equations S6–S9 was used to fit the experimental data normalised by the machine standard (nAUC). Fitting was performed with the SciPy Python package.^53^ To find a global best fit, Monte-Carlo sampling of initial parameters for data fitting was performed 100-times and the solution with the highest adjusted R^2^ was selected.

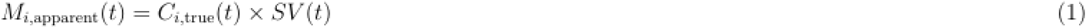

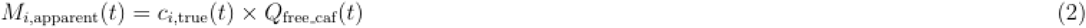

with:

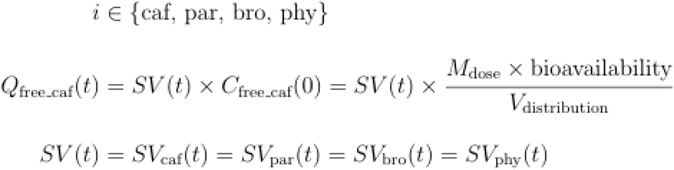

## abbrevations

free,caf: ingested caffeine
caf: absorbed caffeine
par: paraxanthine
bro: theobromine
phy: theophyline

## Acknowledgments

We acknowledge support by the Mass Spectrometry Centre of the Faculty of Chemistry, University of Vienna, and the Joint Metabolome Facility, University of Vienna and Medical University of Vienna. Both facilities are members of the Vienna Life Science Instruments (VLSI). We would also like to thank the Hochschuljubiläumsstiftung (HJS) for their financial support during the course of this research project.

## Additional information

Supplementary Information Supplementary Table

## Author Contributions

J.B. performed research, interpreted data, analysed data and wrote the manuscript, L.N. performed research and analysed data, M.L.F. performed research and analysed data, C.L. performed research and analysed data, M.G. performed research, analysed and interpreted data, B.N. performed research, L.J. performed research and analysed data, A.B. interpreted data and wrote the manuscript, J.Z. analysed data and interpreted data, S.M.M. performed research, analysed and interpreted data and wrote the manuscript, C.G. conceptualized the project, interpreted data and wrote the manuscript.

## Abbreviations

ADME: absorption, distribution, metabolism and excretion
CV: coefficient of variation
FSA: finger sweat analysis
LC: liquid chromatography
MS: mass spectrometry
PCA: principle component analysis
RT: retention time
UHPLC: high performance liquid chromatography

