## Supplementary Information for "Finger Sweat Analysis Enables Short Interval Metabolic Biomonitoring in Humans"

**Table of contents** **Page**

1. Mathematical modelling, equations S2
2. Targeted approach using multiple reaction monitoring (MRM) S3
3. List of supplementary tables S1–S5 S5
4. List of supplementary figures S1–S6 S10

### 1. Mathematical Modelling

#### *Equations*.

$$\frac{\mathrm{d}C_{\mathrm{free\_caf}}}{\mathrm{d}t} = -k_1 C_{\mathrm{free\_caf}} \tag{S1}$$

$$\frac{\mathrm{d}C_{\mathrm{caf}}}{\mathrm{d}t} = k_1 C_{\mathrm{free\_caf}} - k_9 C_{\mathrm{caf}} \tag{S2}$$

$$\frac{\mathrm{d}C_{\mathrm{par}}}{\mathrm{d}t} = k_2 C_{\mathrm{caf}} - k_6 C_{\mathrm{par}} \tag{S3}$$

$$\frac{\mathrm{d}C_{\mathrm{bro}}}{\mathrm{d}t} = k_3 C_{\mathrm{caf}} - k_7 C_{\mathrm{bro}} \tag{S4}$$

$$\frac{\mathrm{d}C_{\mathrm{phy}}}{\mathrm{d}t} = k_4 C_{\mathrm{caf}} - k_8 C_{\mathrm{phy}} \tag{S5}$$

$$c_{\mathrm{caf}}(t) = \frac{C_{\mathrm{caf}}}{C_{\mathrm{free\_caf}}(0)} = \frac{k_1}{k_9 - k_1} \left(e^{-k_1 t} - e^{-k_9 t}\right) \tag{S6}$$

$$c_{\mathrm{par}}(t) = \frac{C_{\mathrm{par}}}{C_{\mathrm{free\_caf}}(0)} = \frac{k_1 k_2}{k_9 - k_1} \left( \frac{e^{-k_9 t}}{k_9 - k_6} - \frac{e^{-k_1 t}}{k_1 - k_6} + \frac{e^{-k_6 t}(k_9 - k_1)}{(k_9 - k_6)(k_1 - k_6)} \right) + \frac{C_{\mathrm{par}}(0)}{C_{\mathrm{free\_caf}}(0)}\, e^{-k_6 t} \tag{S7}$$

$$c_{\mathrm{bro}}(t) = \frac{C_{\mathrm{bro}}}{C_{\mathrm{free\_caf}}(0)} = \frac{k_1 k_3}{k_9 - k_1} \left( \frac{e^{-k_9 t}}{k_9 - k_7} - \frac{e^{-k_1 t}}{k_1 - k_7} + \frac{e^{-k_7 t}(k_9 - k_1)}{(k_9 - k_7)(k_1 - k_7)} \right) + \frac{C_{\mathrm{bro}}(0)}{C_{\mathrm{free\_caf}}(0)}\, e^{-k_7 t} \tag{S8}$$

$$c_{\mathrm{phy}}(t) = \frac{C_{\mathrm{phy}}}{C_{\mathrm{free\_caf}}(0)} = \frac{k_1 k_4}{k_9 - k_1} \left( \frac{e^{-k_9 t}}{k_9 - k_8} - \frac{e^{-k_1 t}}{k_1 - k_8} + \frac{e^{-k_8 t}(k_9 - k_1)}{(k_9 - k_8)(k_1 - k_8)} \right) + \frac{C_{\mathrm{phy}}(0)}{C_{\mathrm{free\_caf}}(0)}\, e^{-k_8 t} \tag{S9}$$

with:

$$C_{\mathrm{free\_caf}} = \frac{M_{\mathrm{dose}} \times \mathrm{bioavailability}}{V_{\mathrm{distribution}}}$$

$$k_9 = k_2 + k_3 + k_4 + k_5$$

$$\mathrm{fc}_i = \frac{k_i}{k_2 + k_3 + k_4} \qquad \text{with} \quad i \in \{2, 3, 4\} \quad \text{for} \quad \{\text{par, bro, phy}\} \tag{S10}$$

abbreviations:

| | | |
|---|---|---|
| free_caf | ... | ingested caffeine |
| caf | ... | absorbed caffeine |
| par | ... | paraxanthine |
| bro | ... | theobromine |
| phy | ... | theophylline |
| fc | ... | fractional conversion |

#### 2. Targeted Approach using multiple reaction monitoring (MRM)

*Chemicals*. Caffeine and formic acid were obtained from Fluka. Caffeine-D9, paraxanthine, theobromine and theophylline were obtained from Sigma-Aldrich. Ultrapure water was obtained from a Millipore system (18.2 MΩ, 185 UV, Millipore), all other solvents were purchased from VWR (LC-MS grade). All chemicals and solvents were used as received.

*Standard Solutions and Calibration Samples*. The stable isotope-labelled caffeine-D9 was used as the internal standard. It was spiked to all samples in a concentration of 10 pg $\mu L^{-1}$. Additionally, a standard mixture containing caffeine, theobromine and theophylline (each 100 pg $\mu L^{-1}$) was measured every thirtieth sample as a quality control. The limit of detection (LOD) was defined as the lowest concentration of the analyte that can be detected with a signal-to-noise ratio of 3 : 1. The lower limit of quantification (LLOQ) was defined as the lowest concentration of the analyte that can be detected with a signal-to-noise ratio of 10 : 1.

*Samples*. The temporal evolution of caffeine, theobromine, theophylline and paraxanthine in fingertips of five volunteers was investigated. This was the suggested number of volunteers by power analysis in order to obtain statistical relevant data (calculated using R-studio with a 10% error rate and a significance criterion of 0.05). Two female and three male volunteers were recruited between 25 and 30 of age. All volunteers were non– to moderate coffee consumers. The volunteers were asked to fast caffeine-containing products for 12 h before the start of each experiment. Subjects presented on 8 am on the study day before the first cup of coffee (equivalent to a double espresso, approx. 80 mg caffeine). Samples from the fingertips were collected before and 1, 3 and 5 h after coffee consumption. The experiment was performed independently on three different days.

*Sample Preparation*. Sample collection was performed similarly as described in the methods section of the main text. The filter paper was transferred into an Eppendorf tube and metabolites were extracted with acetonitrile (500 µL). First, the extraction solution was vortexed for 1 min and then stirred in am Eppendorf Thermomixer comfort 1.5 mL (40 °C, 1400 rpm, 10 min). This process was repeated twice and finally the mixture was centrifuged (10 min, 20’000 rpm). A fraction of the sample solution (250 µL) was transferred into a fresh Eppendorf tube. The samples were dried under a flow of nitrogen. The dried residues were reconstituted in water (250 µL) containing 0.2% formic acid. The extracted samples were sonicated (10 min) and transferred into 96-well plates for analysis.

*Targeted LC-MS/MS*. Targeted LC-MS/MS experiments were performed using a nano- and cap-pump (1260 Infinity, Agilent) together with a microfluidics-based separation system (Chip-Cube, Agilent) hyphenated to a triple quadrupole mass spectrometer (Agilent 6490).

*Liquid Chromatography*. The analyte separation was performed using a chip-based integrated sample enrichment, separation and nanoESI sprayer tip (UHC-CHIP II, ZORBAX 80SB-C18, 5 µm, 25 mm × 75µm enrichment column and 150 mm × 75 µm separation column, Agilent). The injection volume was 0.5 µL. The autosampler was held at 4 °C. Mobile phase A was aqueous solution (0.2% FA) and mobile phase B was acetonitrile (0.2% FA). The gradient included a total run time of 25 min using a flow rate of 400 nL $min^{-1}$. A stepped-gradient was applied by starting with 0% B and increasing to 8% B within 0.1 min. The mobile phase B was linearly increased to 20% (3 min) and then to 80% (5 min), which was held constant for 4 min. Then, the mobile phase B was decreased to 0% and the system allowed to equilibrate for further 16 min. A mixture of isopropanol, acetonitrile, methanol and water (1 : 1 : 1 : 1) was used for needle washes.

*Mass Spectrometry*. The analytes were detected *via* multiple reaction monitoring (MRM) of three different transitions per molecule with a cycle time of 0.8 s and a dwell time of 50 ms per transition (Table S1). Typical MS parameters were as follows: capillary voltage –1.7 to –1.9 kV, gas flow 13 L $min^{-1}$, dry gas temperature 200 °C. Experiments were performed and evaluated using Mass Hunter B.06.00 (Agilent).

#### 3. List of supplementary tables S1–S5

**Table S1**. Caffeine metabolites detected in finger sweat and verified with external analytical standards. n.d. = not detected.

| Compound | Chemical Formula | Monoisotopic mass | pos Precursor $[M+H]^+$ | neg Precursor $[M-H]^-$ | mass error [ppm] | RT [min] |
|---|---|---|---|---|---|---|
| **1,3,7-Trimethyluric acid** | C8H10N4O3 | 210,0753 | n.d. | 209,0680 | 0,96 | 2,86 |
| **1,7-Dimethyluric acid** | C7H8N4O3 | 196,0596 | n.d. | 195,0524 | 0,00 | 2,44 |
| **1-Methyluric Acid** | C6H6N4O3 | 182,0440 | n.d. | n.d. | n.d. | n.d. |
| **1-Methylxanthine** | C6H6N4O2 | 166,0491 | 167,0564 | 165,0413 | 0,60 | 1,75 |
| **3,7-Dimethyluric acid** | C7H8N4O3 | 196,0596 | n.d. | 195,0524 | 0,00 | 2,06 |
| **3-Methyluric acid** | C6H6N4O3 | 182,0440 | n.d. | 181,0367 | 0,55 | 1,52 |
| **3-Methylxanthine** | C6H6N4O2 | 166,0491 | 167,0564 | 165,0413 | 0,00 | 1,60 |
| **5-Acetylamino-6-formylamino-3-methyluracil** | C8H10N4O4 | 226,0702 | 227,0775 | n.d. | 0,44 | 1,09 |
| **7-Methyluric acid** | C6H6N4O3 | 182,0440 | n.d. | 181,0367 | 0,55 | 1,34 |
| **7-Methylxanthine** | C6H6N4O2 | 166,0491 | 167,0564 | 165,0413 | 1,79 | 1,48 |
| **Caffeine** | C8H10N4O2 | 194,0804 | 195,0877 | n.d. | 0,51 | 3,28 |
| **Paraxanthine** | C7H8N4O2 | 180,0647 | 181,0720 | n.d. | 0,00 | 2,55 |
| **Theobromine** | C7H8N4O2 | 180,0647 | 181,0720 | n.d. | 0,55 | 2,14 |
| **Theophylline** | C7H8N4O2 | 180,0647 | 181,0720 | n.d. | 1,10 | 2,65 |
| **Uric Acid** | C5H4N4O3 | 168,0283 | 169,0356 | 167,0205 | 1,18 | 0,78 |
| **Xanthine** | C5H4N4O2 | 152,0334 | 153,0407 | 151,0256 | 1,31 | 0,95 |

**Table S2**. Recovered parameters. Kinetic parameters [$h^{-1}$], initial concentrations and caffeine amounts (Qfree_caf) [nAUC] fitted by the kinetic model (Figure 5A). As comparison the kinetic parameters found in blood plasma after uptake of 5 mg caffeine per kg specimen body mass are listed (Bonati 1982).

| Parameter | Donor 1 | Donor 2 | Bonati et al. |
|---|---|---|---|
| $k_1$ | 0.214 | 0.179 | 6.31 ± 1.91 |
| $k_2$ | 0.082 | 0.052 | 0.099 ± 0.022 |
| $k_3$ | 0.027 | 0.375 | 0.013 ± 0.001 |
| $k_4$ | 0.016 | 0.007 | 0.015 ± 0.002 |
| $k_5$ | 0.077 | 0.000 | - |
| $k_6$ | 0.382 | 0.302 | - |
| $k_7$ | 0.266 | 0.582 | - |
| $c_{paraxanthine}(0)$ | 0.002 | 0.000 | - |
| $c_{theobromine}(0)$ | 0.002 | 0.001 | - |
| $c_{theophylline}(0)$ | 0.000 | 0.000 | - |
| $Q_{free\_caf}(0)$ | 214 | 910 | - |
| $Q_{free\_caf}(15)$ | 1000 | 735 | - |
| $Q_{free\_caf}(30)$ | 347 | 271 | - |
| $Q_{free\_caf}(45)$ | 263 | 160 | - |
| $Q_{free\_caf}(60)$ | 247 | 88 | - |
| $Q_{free\_caf}(90)$ | 137 | 63 | - |
| $Q_{free\_caf}(120)$ | 127 | 13 | - |
| $Q_{free\_caf}(180)$ | 97 | 34 | - |
| $Q_{free\_caf}(240)$ | 96 | 60 | - |
| $Q_{free\_caf}(360)$ | 90 | 34 | - |
| $Q_{free\_caf}(480)$ | 120 | 98 | - |
| $Q_{free\_caf}(1440)$ | 329 | 710 | - |
| $Q_{free\_caf}(1500)$ | 224 | 901 | - |
| $Q_{free\_caf}(1560)$ | 293 | 516 | - |
| $Q_{free\_caf}(1620)$ | 1000 | 1000 | - |

**Table S3**. Fractional conversion of caffeine to degradation products. The fractional conversions of caffeine to degradation products (paraxanthine, theobromine, theophylline) in % for donors 1 and 2 and a comparison to literature are given. Values from donors 1 and 2 and Bonati et al. were calculated with Equation S10.

| Degradation Product | Donor 1 | Donor 2 | Bonati et al. | Lelo et al. |
|---|---|---|---|---|
| **Paraxanthine** | 66 | 12 | 78 ± 17 | 84 ± 5 |
| **Theobromine** | 22 | 86 | 12 ± 1 | 12 ± 4 |
| **Theophylline** | 13 | 2 | 10 ± 2 | 4 ± 1 |

**Table S4**. Parameters for the targeted LC-MS/MS approach.

| Analyte | Precursor Ion | Product Ion | Dwell-time (ms) | Fragmentor/Collision Energy (eV) |
|---|---|---|---|---|
| **Caffeine** | 195.1 | 138 | 50 | 380/40 (quantifier) |
| | | 110 | 50 | 380/40 |
| | | 83 | 50 | 380/40 |
| **Theophylline/ Paraxanthine** | 181 | 123.9 | 50 | 380/30 |
| | | 95.9 | 50 | 380/30 |
| | | 69 | 50 | 380/30 (quantifier) |
| **Theobromine** | 181 | 122.2 | 50 | 380/30 |
| | | 107.9 | 50 | 380/30 |
| | | 67 | 50 | 380/30 (quantifier) |

**Table S5**. Analytical validation of CF and its metabolites in the proof-of-principle study: The compounds were spiked in two concentrations. The precision for CF, TB and TP is between 0.03 and 15.6% with overall process efficiency from 88–92%, including the associated coefficients of variation (CV) in brackets. The LLOQ for caffeine and its metabolites was defined as the lowest concentration giving signal-to-noise ratio of at least 10. The lowest concentration that can be detected, with a signal-to-noise ratio of 3:1, is specified as the limit of detection (LOD).

| Compound | Spiked conc. [pg/FP] | Precision [%] | Overall Process Efficiency (CV) | LLOQ [pg/FP] | LOD [pg/FP] |
|---|---|---|---|---|---|
| **Caffeine** | 0.5<br>50 | 15.9<br>0.03 | -<br>88.4 (7.2%) | 0.54 | 0.22 |
| **Theobromine** | 0.5<br>50 | 4.1<br>0.7 | -<br>92.0 (5.9%) | 0.68 | 0.28 |
| **Theophylline** | 0. 5<br>50 | 1.9<br>0.2 | -<br>89.9 (7.5%) | 0.42 | 0.20 |

#### 4. List of supplementary figures S1–S6

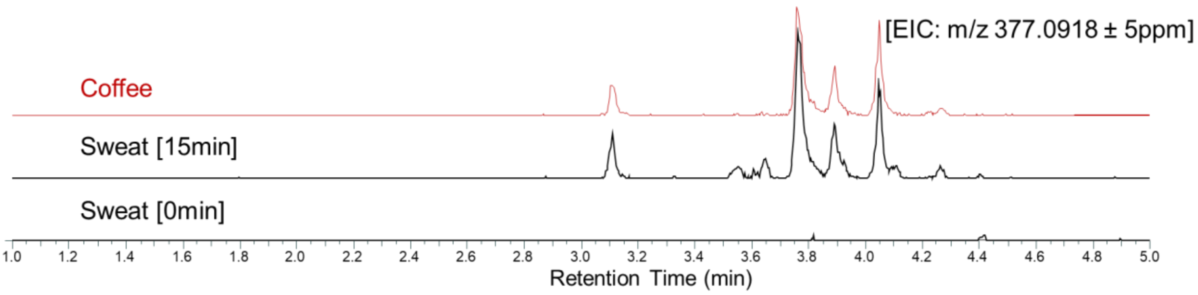


**Supplementary Figure S1**. Similarity of extracted ion chromatograms (EIC) of the unidentified feature 377.0918 and its isomers regarding the source (coffee) and the detection in finger sweat samples 15 minutes (15min) after coffee consumption is shown. The corresponding finger sweat sample collected just before coffee consumption (0min) served as negative control.

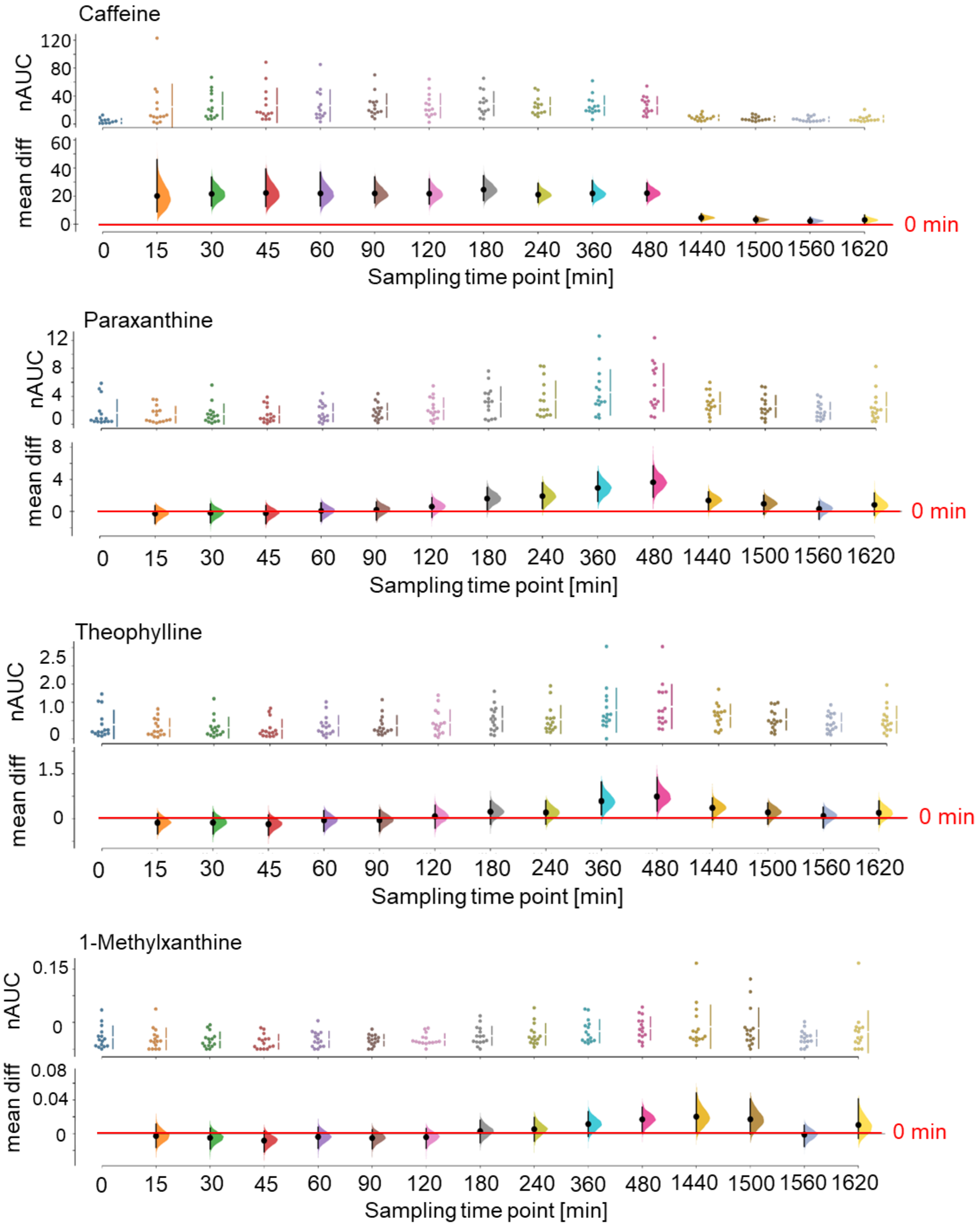


**Supplementary Figure S2**. Effect size plots with data from 14 individuals for caffeine, paraxanthine, theophylline and 1-methylxanthine are shown. nAUC, normalized area under curve; mean diff, mean difference.

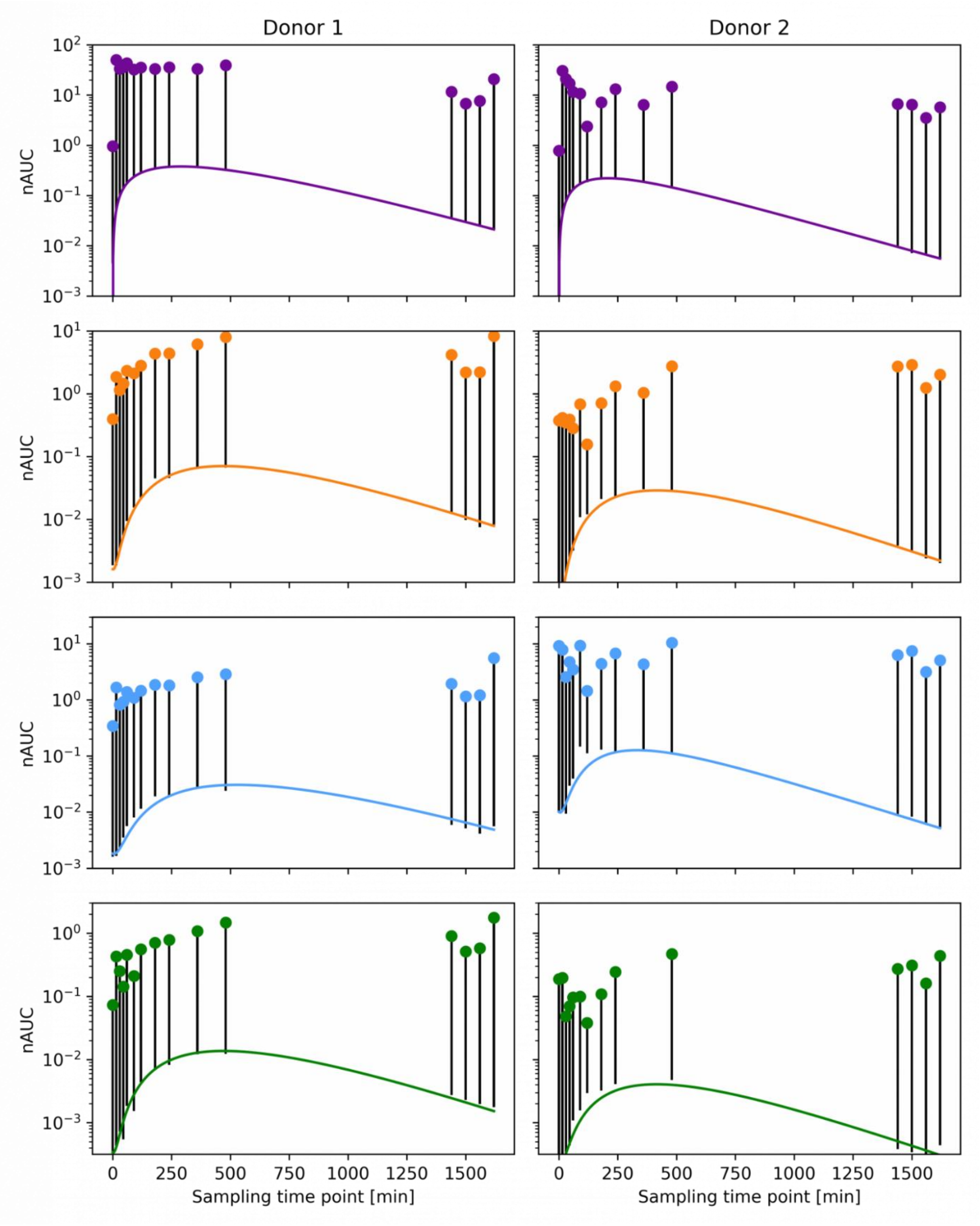


**Supplementary Figure S3**. Visual representation of the influence of $Q_{free\_caf}(t)$. Normalized concentration time series of caffeine, paraxanthine, theobromine and theophylline (first to last row in that order) for donor 1 and donor 2 (left and right column respectively). The symbols refer to the measured values, the coloured lines refer to the fitted values and the black bars indicate how much of the difference is described by $Q_{free\_caf}(t)$. Note that the y-axis is logarithmic.

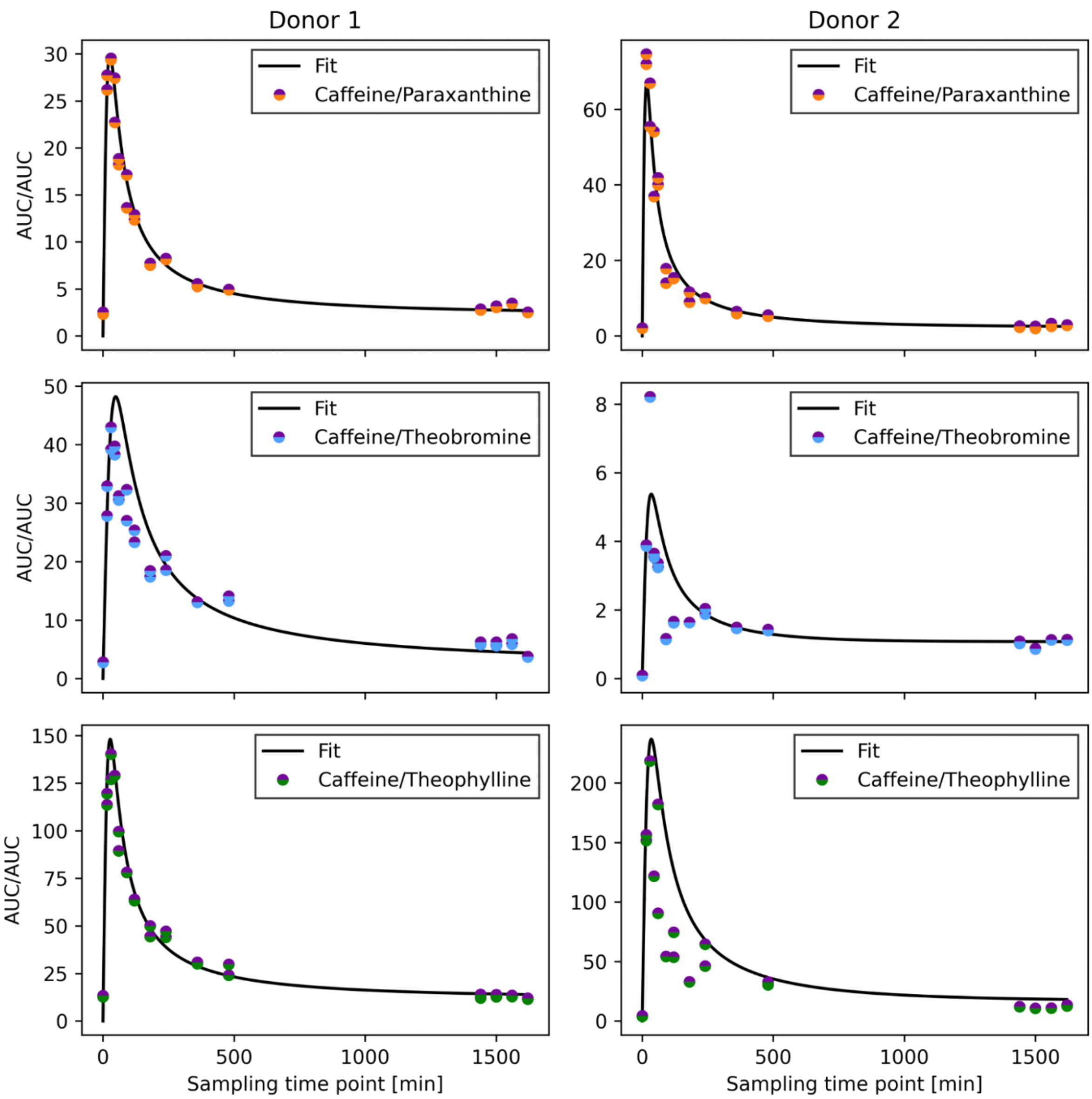


**Supplementary Figure S4**. The sweat volume cancels out upon division of two metabolites. The noise introduced by the different sweat volumes cancels out upon division of the concentration time series of two metabolites as shown here for caffeine and its major degradation products for donor 1 and 2 (left and right column respectively). The fit curve corresponds to the division of the fitted time-curves of the respective metabolite pair.

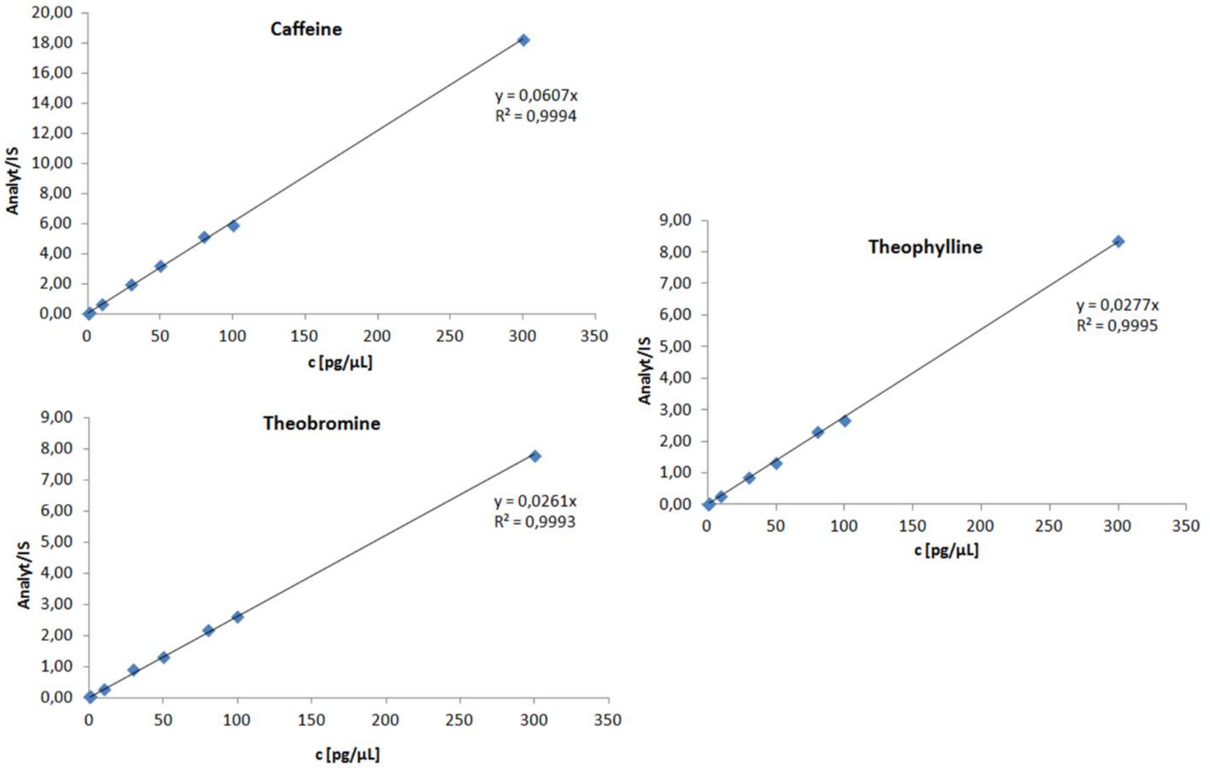


**Supplementary Figure S5**. Calibration curves for caffeine, theobromine and theophylline in eight concentration levels in the range of 0.5–300 pg/µL (0.25–150 pg/µL on column) obtained using the microfluidics-based chip-cube separation system coupled to a triple quadrupole mass spectrometer with their corresponding correlation factors ($R^2$) are shown. IS, internal standard.

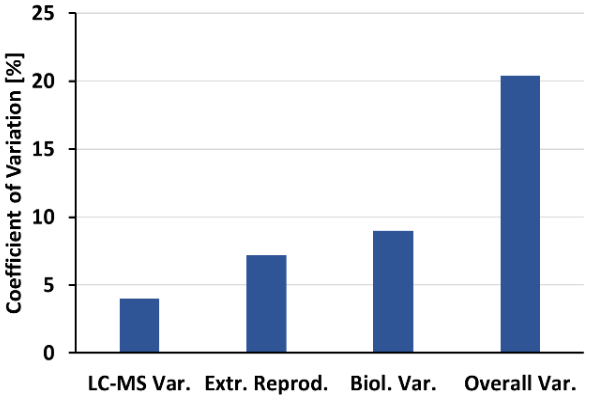


**Supplementary Figure S6**. Coefficients of variation of caffeine quantification from the fingertips using the microfluidic-based chip-cube separation system coupled to a triple quadrupole mass spectrometer. The LC-MS variation represents the coefficients of variations of 3 technical replicates. The extraction reproducibility represents the coefficient of variation of 3 biological extractions, each with 3 technical replicates. The biological variation indicates the coefficients of variation of the average caffeine amount after 5 h of coffee intake of all the donors. The overall variation represents the coefficients of variations of all technical and biological replicates of 5 donors after 5 h of coffee intake.
